# Computational modeling of seizure spread on a cortical surface

**DOI:** 10.1101/2020.12.22.423957

**Authors:** Viktor Sip, Maxime Guye, Fabrice Bartolomei, Viktor Jirsa

## Abstract

In the field of computational epilepsy, neural field models helped to understand some large-scale features of seizure dynamics. These insights however remain on general levels, without translation to the clinical settings via personalization of the model with the patient-specific structure. In particular, a link was suggested between epileptic seizures spreading across the cortical surface and the so-called theta-alpha activity (TAA) pattern seen on intracranial electrographic signals, yet this link was not demonstrated on a patient-specific level. Here we present a single patient computational study linking the seizure spreading across the patient-specific cortical surface with a specific instance of the TAA pattern recorded in the patient. Using the realistic geometry of the cortical surface we perform the simulations of seizure dynamics in The Virtual Brain platform, and we show that the simulated electrographic signals qualitatively agree with the recorded signals. Furthermore, the comparison with the simulations performed on surrogate surfaces reveals that the best quantitative fit is obtained for the real surface. The work illustrates how the patient-specific cortical geometry can be utilized in The Virtual Brain for personalized model building, and the importance of such approach.

## 1. Introduction

In the field of epilepsy, computational modeling is used to both understand the general principles of seizure initiation, progression, and termination, as well as to analyze and interpret the clinical data for individual patients. These can include non-invasive imaging revealing the geometry and connectivity of the patient’s brain as well as invasive intracranial EEG recordings, both ictal and interictal. Recent studies indicate that some features of the large-scale organization of epileptic seizures may be better understood through investigation of the phenomena acting on smaller scales than the coarse parcellation of the brain into tens of discrete brain regions which is often employed in whole-brain models of epilepsy (Taylor et al., 2014; Jirsa et al., 2017). As two examples, Proix et al. (2018) introduced a neural field model of epilepsy dynamics that gives rise to the slow propagation of the ictal wavefront, fast propagating waves in recruited regions, and clustered synchronous termination of the seizures. Sip et al. (2020) suggested a link between the slow seizure propagation across the cortex with the so-called theta-alpha activity onset pattern visible on intracranial EEG signals. Such phenomena are strongly linked to the continuous nature of the underlying substrate and cannot be captured with the network-based models which lack the spatial component.

In the recent years, more research aims to personalize the large-scale network models in order to provide insight or predictions at individual level. This personalization can take many forms, such as using the patient-specific structural connectome derived from diffusion-weighted imaging (Jirsa et al., 2017), using the functional connectome obtained from intracranial recordings (Goodfellow et al., 2016; Sinha et al., 2017), or estimating the model parameters from recorded seizures (Hashemi et al., 2020). However, exploiting the individual data in the models operating at smaller scales to gain clinically relevant insight is rare, as they tend to explore general phenomena of epileptic seizures and not their specific instances.

In this work we demonstrate how such personalization of a neural field model can be performed using The Virtual Brain (TVB) platform (Sanz Leon et al., 2013). TVB is a simulator of brain dynamics, easily allowing to integrate patient-specific structural data. It supports both network based modeling, where the brain is represented as a set of connected nodes, as well as surface based modeling, where the cortex is represented as a triangulated surface, and the nodes on the surface are connected to each other both locally and through long-range connections. In the context of epilepsy, however, only the former approach was utilized so far (Jirsa et al., 2017).

We present a single-patient study that in TVB links the patient-specific cortical surface with the patient-specific stereo-electroencephalographic (SEEG) recording of an epileptic seizure. Specifically we investigate the theta-alpha activity pattern (TAA, Fig. 1), characterized by sustained oscillations in the *θ* − *α* range with gradually increasing amplitude (Alarcon et al., 1995; Perucca et al., 2014; Lagarde et al., 2016). The pattern was reported to occur in 6-23% patients (Singh et al., 2015) and it was identified as the most common pattern in the areas of seizure propagation (Perucca et al., 2014). In the previous work we suggested that the pattern is caused by spreading seizures across the cortical surface (Sip et al., 2020, Fig. 2A). Crucial to our presented perspective is the distinction between the source (i.e. the gray matter tissue) and sensor (i.e. SEEG) spaces and the link between them, defined by the sensor-to-source projection. We hypothesize that seizure spread along the folded cortical surface may mimick dynamic spatiotemporal characteristics of seizure classes in sensor space that are not present in source space and thus lead to misinterpretation of the spatiotemporal organization of the seizure.

**Figure 1:**
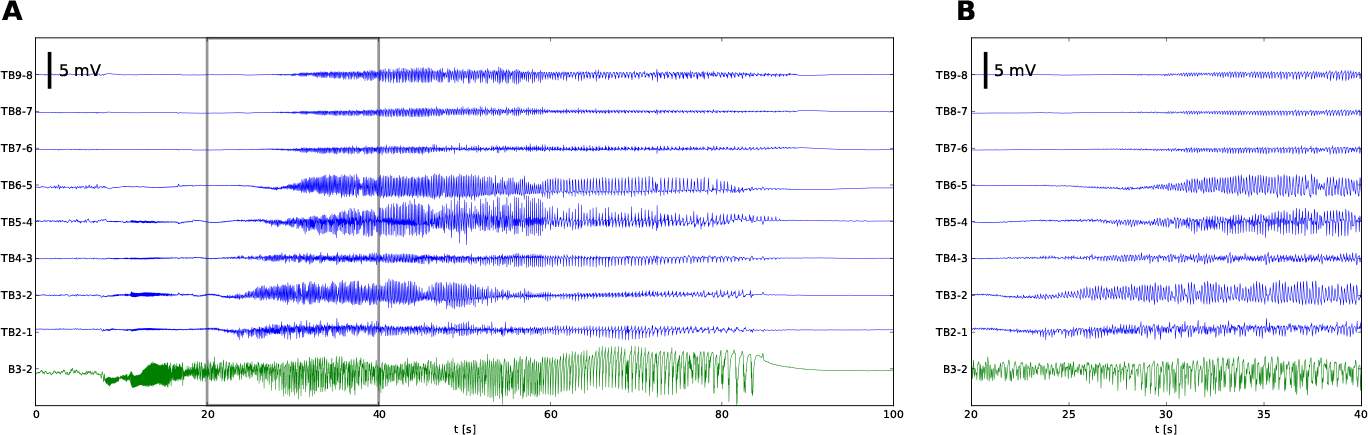
TAA pattern recorded in a patient with temporal lobe epilepsy. (A)Stereo-electroencephalographic (SEEG) traces of the recorded seizure in bipolar representation. (B) Detail of the seizure onset, delimited by the gray lines in (A). The onset of ictal activity is first recorded on B3-2 (in green) located in the right hippocampus. The seizure then spreads to the TB contacts (in blue) located in the right temporal lobe. There the onset shows the distinguishing features of the TAA pattern: oscillating activity with frequency of 8 Hz and slowly increasing amplitude. It is worth noting that at the time of the initial oscillations in the right hippocampus (8 - 18 s), some oscillatory activity appears also on the mesial contacts of TB electrode, but soon disappears. We assume that this initial activity visible on TB electrode is due to the volume conduction from the hippocampus, and does not reflect the source activity close to the TB contacts (see Fig. S1 for details), and we exclude this initial period from the analysis presented later.

**Figure 2:**
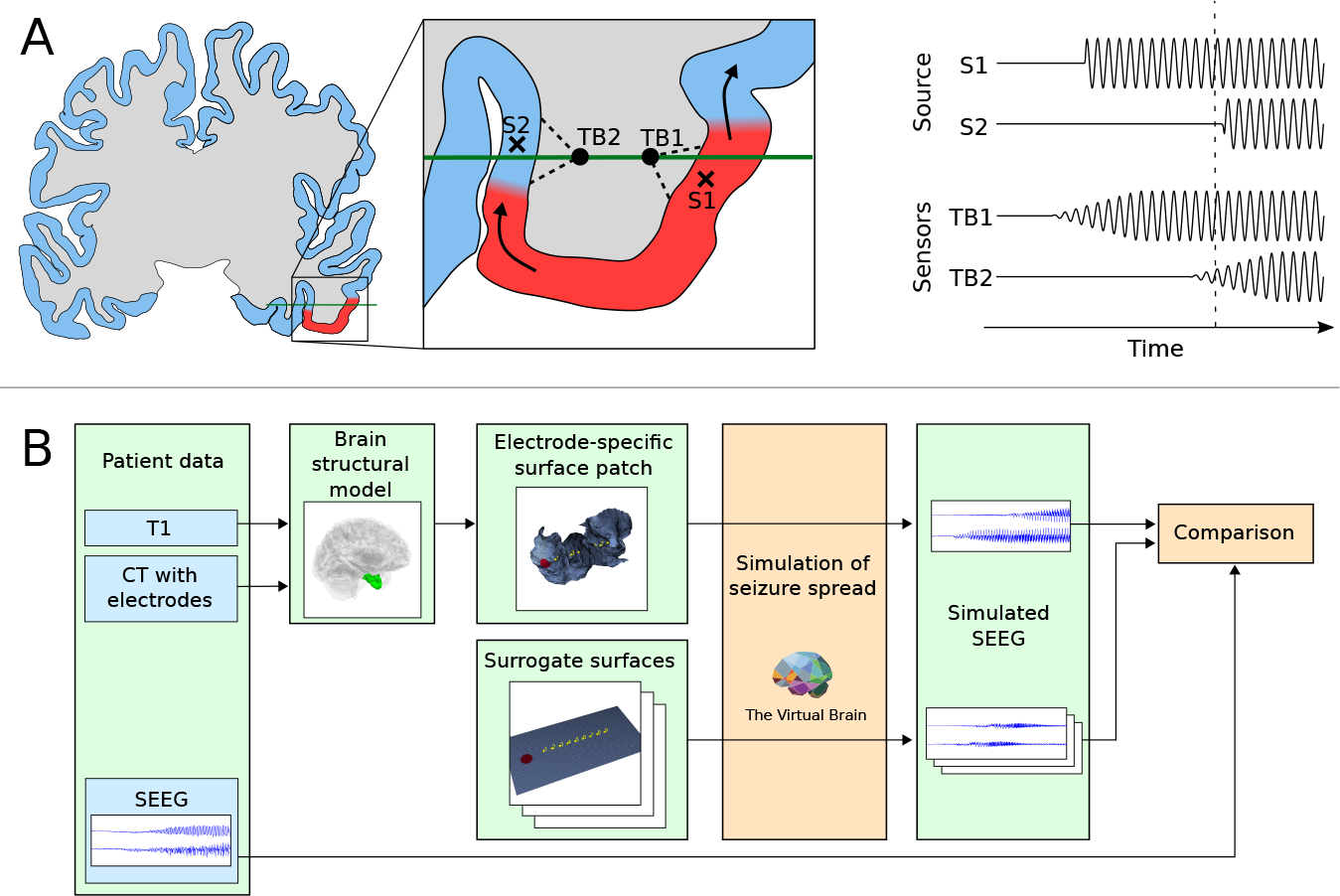
(A) Hypothesized mechanism behind the TAA pattern, for clarity shown schematically on a two-dimensional brain slice. Left: The seizure spreads across the cortical surface and recruits the cortical tissue from the normal (blue) to the seizing state (red). The generated local field potentials are measured by the contacts (TB1, TB2) of the implanted electrode (green). Right: Time series of the source activity and recorded signals from the SEEG sensors. Dashed vertical line indicates the time of the snapshot in panel A. Every unit (S1, S2) on the cortical surface enters the seizure state through a rapid transition. Due to the spatial averaging of the source activity performed by the SEEG sensors (via the LFP summation), the recorded seizure onset on the sensors (TB1, TB2) is gradual with slowly increasing amplitude of the oscillations. (B) Outline of the workflow. From patient’s imaging data the structural model of the brain is built and the surface patch around the electrode of interest is extracted. On this and three surrogate surfaces the simulations of seizure spread are performed, and simulated SEEG signals are compared qualitatively and quantitatively with the recorded SEEG.

To illustrate this point, we demonstrate that this scenario is plausible on the spatial and temporal scales relevant to epileptic seizures in human brain by means of numerical simulation (Fig. 2B). We select a single epileptic patient for whom the TAA pattern was observed and extract a patch of the cortical surface around the electrode where it was seen. We then simulate the seizure spread on this surface using TVB. As a mathematical model we use the Epileptor (Jirsa et al., 2014), a phenomenological model of seizure dynamics, in its field formulation (Proix et al., 2018). We then compute the forward solution from source to sensor space and examine whether the features of the electrographic pattern on the simulated SEEG are qualitatively consistent with the real SEEG recordings for the specific patient. Furthermore, we perform the simulations also on surrogate surfaces to demonstrate that the best quantitative fit is obtained for patient-specific surface.

## 2. Methods

### 2.1. Patient data

From the cohort of 15 epileptic patient in the previous study (Proix et al., 2017), we selected a single patient and a single electrode with the clearest TAA pattern present on more than eight contacts of the electrode. The selected patient was a 29-year-old male with temporal lobe epilepsy. The epileptogenic zone was estimated by the clinicians to be in the right hippocampus, right fusiform gyrus, right entorhinal cortex, and right temporal pole. The patient signed an informed consent form according to the rules of local ethics committee (Comité de Protection des Personnes Sud-Méditerranée I).

The patient underwent standard clinical evaluation, described in detail before (Proix et al., 2017). The T1 weighted images (MPRAGE sequence, repetition time = 1900 ms, echo time = 2.19 ms, 1.0 × 1.0 × 1.0 mm, 208 slices) were obtained on a Siemens Magnetom Verio 3T MR-scanner. As a part of the clinical evaluation, the patient was implanted with 13 stereotactic EEG electrodes (10 right hemisphere, 3 left hemisphere). Each electrode has 10 to 15 contacts (2 mm long a 0.8 mm in diameter), which are separated by 1.5 mm from each other. The SEEG was recorded by a 128 channel Deltamed™ system using a 256 Hz sampling rate. The recordings were band-pass filtered between 0.16 and 97 Hz by a hardware filter. After the electrode implantation, a CT scan of the patient’s brain was acquired to obtain the location of the implanted electrodes.

### 2.2. Extracting the cortical surface

The brain anatomy was reconstructed from the T1 weighted images by FreeSurfer v6.0.0 (Fischl, 2012) using the *recon-all* procedure. The CT scan with the implanted electrodes was aligned with the T1 weighted images using the linear registration tool FLIRT from the FSL toolbox (Jenkinson et al., 2002). The position of the contacts was then determined by reading the position of the innermost and outermost contact in the MRtrix image viewer (Tournier et al., 2012), and placing the other contacts on this line using the known spacing of 3.5 mm between the contacts.

The part of the cortical surface used in this study was then obtained by the following steps: First, we took a midsurface of the pial surface and white matter-gray matter interface, i.e. the surface lying halfway between these surfaces. Then, the parts of the surface further than 15 mm from any contact on the selected electrode were discarded. After that, only the largest connected part of the remaining surface was kept and the small components unconnected to the largest were discarded as well.

To obtain a triangular mesh fine enough to properly resolve the spatiotemporal dynamics of the simulated seizure spread, the extracted part of the triangulated surface generated by FreeSurfer was further refined. This was done by splitting every triangle of the original triangulation into four triangles with the vertices located at the vertices of the original triangle and at the midpoints of the original triangle’s edges.

### 2.3. Epileptor field model

In this study the Epileptor model (Jirsa et al., 2014) was employed in its field formulation (Proix et al., 2018). This phenomenological neural mass model was originally developed to produce the dynamics observed in *in vitro* epileptic tissue. This was done by identifying the invariant features of the seizure-like activity and classifying them as bifurcations of the underlying dynamical system. The created model was mathematically represented by a set of five integro-differential equation, representing a fast population (*u*_1_ and *u*_2_ variables) generating rapid discharges, a population acting on an intermediate time scale (*q*_1_ and *q*_2_) generating slower oscillations, and a slow variable *v* guiding the system into and out of the seizure states. As a phenomenological model, the Epileptor does not provide a one-to-one mapping between its variables and the observables of the electrochemical system of the actual brain, although the authors discussed how the variables of the model can be represented in such systems.

In the field formulation employed, the dynamics are defined on a spatially continuous sheet (or line in one dimension), and any two points on the sheet are coupled through a local connections of decreasing strength with increasing distance. This field model was proved to be capable of producing a slowly progressing seizure front, traveling waves as well as a simultaneous termination of a seizure in distant areas (Proix et al., 2018). The variables of the model are functions of space ***x*** and time *t*; where not necessary, we omit explicitly writing these dependencies for brevity. The model is described by five integro-differential equations,

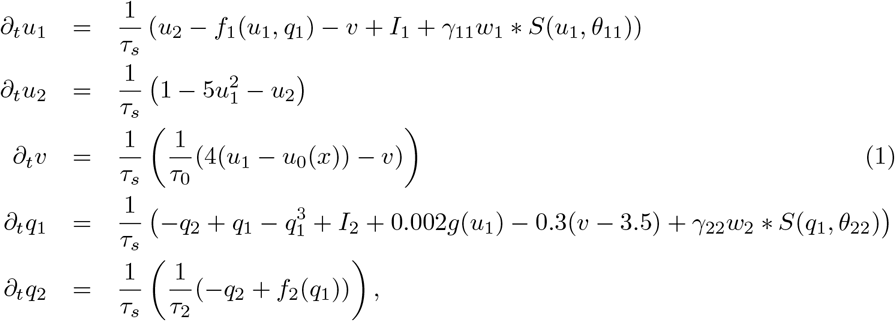

where

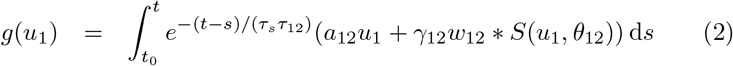

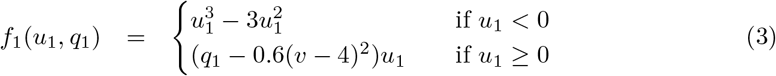

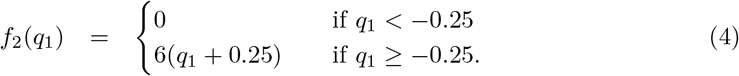

The local connections in space are expressed via the convolution operator,

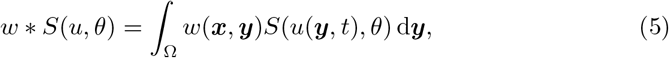

where the local activity *u* is passed through a Heaviside function *S*(*u, θ*) = *H*(*u* − *θ*). The Laplacian local connection kernel is used, 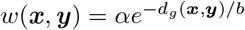 with *d*_*g*_(***x, y***) being the geodesic distance between the two points ***x*** and ***y*** on the surface, and with a normalization constant 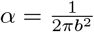.

Following parameter values were used for all presented simulations: *b* = −1, *θ*_11_ = −1, *θ*_12_ = −1, *θ*_22_ = −0.5, *γ*_12_ = −10, *γ*_22_ = −1, *I*_1_ = 3.1, *I*_2_ = 0.45, *τ*_2_ = 100, *a*_12_ = 3. Rest of the parameters differed in the simulations presented and their values are given in Tab. 1.

**Table 1:** Simulation parameters. *τ*_0_, *τ*_*s*_ - temporal constants of the Epileptor model, *γ*_11_ - coupling strength of the fast subsystem, 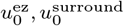 - epileptogenicity of the epileptogenic zone and of the surrounding tissue, Δ_*t*_ - time step of the numerical integration. Symbol * marks a variable parameter. Values of *γ*_11_ for cortical sheet simulations on different surfaces are given in Tab. 2.

| Parameter | $\tau_0$ | $\tau_s$ | $\gamma_{11}$ | $u_0^{\text{ez}}$ | $u_0^{\text{surround}}$ | $\Delta_t$ |
| --- | --- | --- | --- | --- | --- | --- |
| Value | 20000 | 5.88 | * | -1.8 | -2.3 | 0.2 |

With the parameters used in the paper, an unconnected Epileptor exhibits different behavior based on the value of the excitability parameter *u*_0_. For 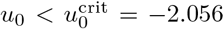, the system converges to a stable fixed point, considered to represent the normal, healthy regime. In this fixed point, the values of the slower variables *q*_1_, *q*_2_, and *s* place the fast subsystem (*u*_1_, *u*_2_) in the monostable state for 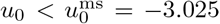, and in the bistable state for 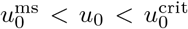. For 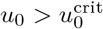 the system periodically switches between a silent and oscillatory state.

In the previous works (Jirsa et al., 2014; Proix et al., 2018) the oscillations generated by the subsystem *q*_1_, *q*_2_ were identified with Spike-and-Wave Discharges (SWD). Since here we use the subsystem to generate oscillations with frequencies above the generally accepted range of 2-4 Hz of SWDs, we avoid this terminology in order not to cause any confusion. Instead we call the *q*_1_, *q*_2_ subsystem simply an *intermediate subsystem* of the Epileptor and we call the traveling waves supported by this system *fast waves*, referring to the high velocity of such waves compared to the velocity of seizure spread.

### 2.4. Numerical simulations

The numerical simulations of the system (1) on the triangulated surface were performed in The Virtual Brain (TVB) platform (Sanz Leon et al., 2013) in version 1.5.4 with some modifications specific to surface-based simulations. For all simulations, the field was initially placed in the stable fixed point of the unconnected Epileptor with the excitability value 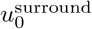, and the temporal integration was then performed using Heun’s method with time step Δ_*t*_ given in Tab. 1.

#### Boundary conditions

There were no local connections from outside of the modeled part of the cortical surface. With the employed coupling, this can be also interpreted as modeling the whole cortex and keeping the Epileptors outside of the modeled domain in a stable state so that the coupling terms *S*(*u*_1_, *θ*_11_), *S*(*u*_1_, *θ*_12_) and *S*(*u*_2_, *θ*_22_) are all zero. This could be achieved by setting the excitability parameter *u*_0_ of the outside tissue to be very low, so that a seizing tissue in the modeled part would not excite it.

### 2.5. Forward model for SEEG signals

In the human cortex, the most numerous neuron type is the pyramidal cell. Due to their geometrical structure with long dendrites oriented perpendicularly to the cortical surface, they might be well represented as electrical dipoles (Buzsáki et al., 2012). Following this idea, we assume that each point on the cortical surface acts as electrical dipole. In the spatially continuous formulation the local field potential measured by the electrode contact at point ***x***_*s*_ generated by the source activity *s*(***x***, *t*) on the surface Ω is given by

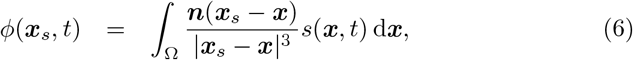

where ***n*** is the outward oriented normal of the surface. In the discretized version using the calculated solution on a triangulated surface this reads

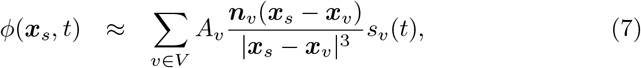

where *V* is the set of all vertices on the triangulated surface, *A*_*v*_ is the area associated with a vertex (calculated as one third of the sum of areas of neighboring triangles), ***n***_*v*_ is the outwards oriented normal of a vertex (calculated as a weighted average of the normals of neighboring triangles), ***x***_*v*_ is the position of the vertex, and *s*_*v*_(*t*) is the calculated activity at the vertex. In the implementation, the temporally independent part of right hand side of Eq. (7) is precomputed and stored in the gain matrix *G* of size *n*_sensors_ × *n*_vertices_. The calculation of the measured signal can then be written in a vector form,

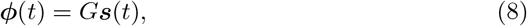

where ***ϕ*** is the vector of signals at given locations, and ***s*** is the vector of simulated activity at all vertices. Following the original Epileptor paper, we identify the source activity with the difference of two state variables of Epileptor model, *s*(*t*) = *q*_1_(*t*) − *u*_1_(*t*) (Jirsa et al., 2014).

### 2.6. Analysis of the TAA onset timing

To provide a quantitative comparison of the timing of the TAA pattern appearance in the recorded and simulated SEEG signals we determined the TAA onset time for all bipolar signals. By *onset time* we mean the time point where the amplitude of the signal starts to grow. It was determined as the time when the signal envelope rises above 20% of its maximum value obtained during the seizure. To calculate the envelope the signals were first high-pass filtered to remove the low frequency oscillations (third order Butterworth filter, cutoff frequency 0.2 Hz). Next, the envelope of the signal was calculated by rectifying the signal (i.e. taking its absolute value) and low-pass filtering the result (third order Butterworth filter, cutoff frequency 0.6 Hz). The calculated onset times were shifted so that the earliest onset time is at zero.

### 2.7. Seizure spread velocity in the model

We determined the dependence of the seizure spread velocity on the coupling strength *γ*_11_ in the model via numerical simulations. We performed multiple numerical simulations of the seizure spread on a rectangular cortical sheet of dimensions 40 × 20 mm with different values of *γ*_11_. On the narrow side of the domain an epileptogenic zone of width 10 mm was placed. The parameters of the simulations and the values of the epileptogenicity of the epileptogenic zone and of the surrounding tissue were the same as in the simulation of the folded cortical surface (Tab. 1). Two points on the domain midline were selected (10 and 30 mm from the edge of the epileptogenic zone) and the seizure spread velocity in each simulation was calculated as the fraction of the distance between the two points and the temporal difference of the simulated seizure onset in these points.

### 2.8. Estimation of the fast wave velocity in the model

The velocity of the fast waves in the simulations was estimated as follows. We selected a time point (*t* = 20 s) after the whole patch was recruited and based on a visual inspection of the simulation results we noted that the fast waves in all simulations travel in the lateral-mesial direction. We then found a shortest path on the triangulated surface connecting the vertices closest to the innermost and outermost contact of the implanted electrode and selected 30 vertices located uniformly on this path. Next, we took a 1 s long time series of the *q*_1_ variable in these vertices after the selected time point. For all values of the fast wave velocities *u*_fast_ in the range from 50 mm s^−1^ to 1000 mm s^−1^ discretized by 200 points we shifted the time series by the temporal offset 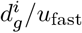 (where 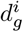 is the geodesic distance of the *i*-th point on the path from the first point), and we selected the value of *u*_fast_ where the average correlation coefficient of all pairs of shifted signals was highest.

## 3. Results

### 3.1. Model of the folded cortical surface

We simulated the seizure propagation on a part of a cortical surface of an epileptic patient. The simulated part of the cortical surface (Fig. 3B, denoted by shorthand ‘Real’) represented the part of the cortical surface located in the vicinity of a single electrode, where the TAA pattern (shown on Fig. 1) was recorded. Parameters of the triangulated patch are given in Tab. 2.

**Figure 3:**
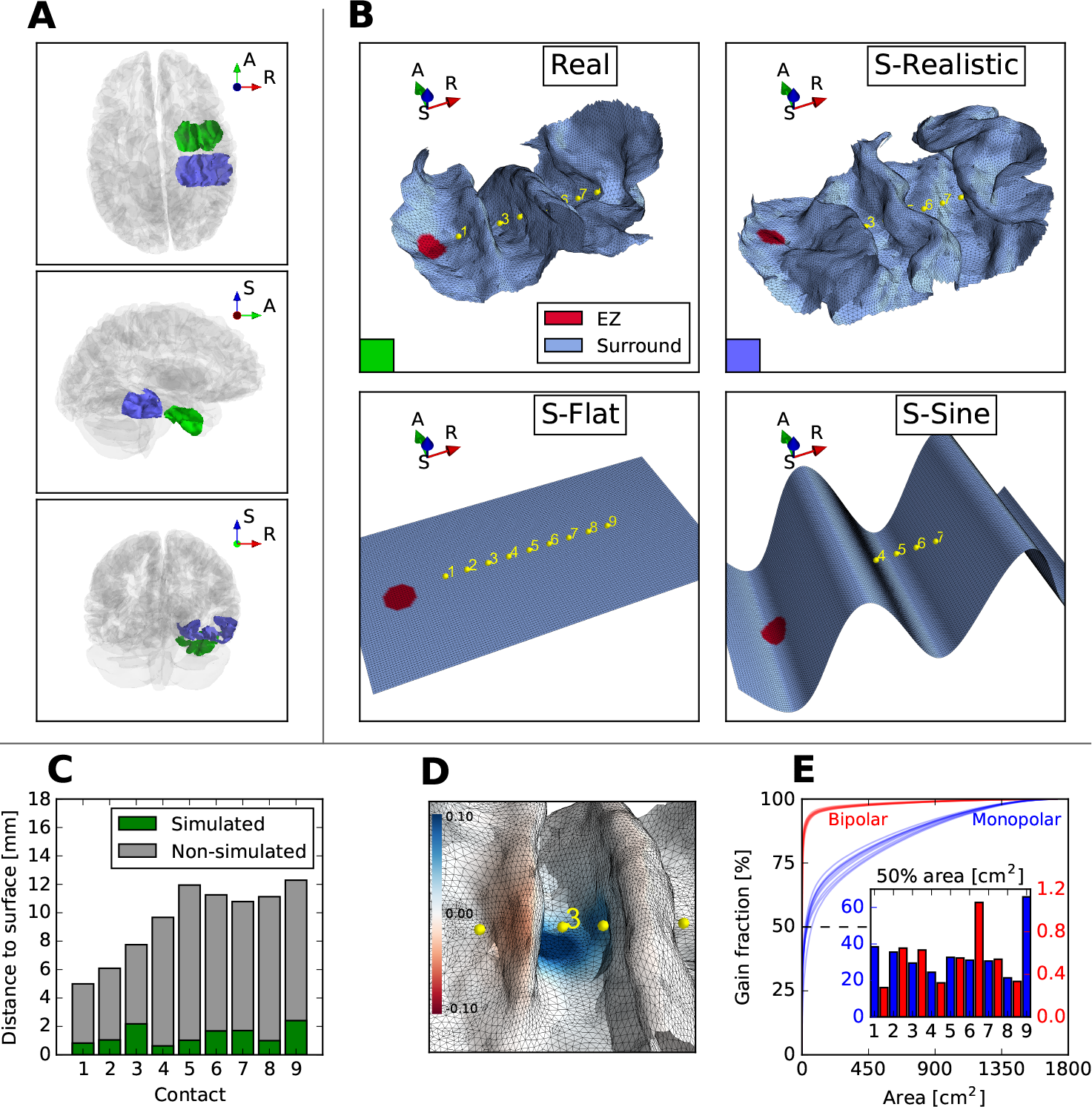
(A) Position of the modeled patches in the whole brain: Real (green) and S-Realistic (blue) surfaces. Axes notation: R - Right, A - Anterior, S - Superior. (B) Detail of the real and three surrogate surfaces. Electrode contacts (in yellow) are numbered from the 1 on the mesial side to 9 on the lateral side. The distance between the neighboring contacts is 3.5 mm. Surface coloring shows the position of the epileptogenic zone (EZ). (C) Minimal distance of every electrode contact from the part of the cortical surfaces included in the simulation (green) and from the subcortical structures or the part of the cortical surface not included in the simulation (gray). (D) Spatial representation of the gain matrix row for contact 3. Blue colors marks the positive weighting, red color negative. (E) Fraction of the gain matrix associated with the area of given size for monopolar (blue) and bipolar (red) referencing. See the main text (Sec. 3.3) for further details. Inset: area associated with 50% of the gain matrix for monopolar (blue) and bipolar (red) referencing. Note the different scales for monopolar and bipolar referencing.

**Table 2:** Parameters of the triangulated surfaces (the real surface and three surrogate surfaces described in Sec. 3.2). Last column shows the coupling strength *γ*_11_ adjusted in the simulations to obtain the same propagation delay between the first and last contact pairs as in the real patient’s recording.

| Surface | $N_{\text{triangles}}$ | $N_{\text{vertices}}$ | Area [mm<sup>2</sup>] | Edge length [mm] | | | $\gamma_{11}$ |
| --- | --- | --- | --- | --- | --- | --- | --- |
|  |  |  |  | Min | Mean | Max |  |
| Real | 24352 | 12424 | 2350.4 | 0.079 | 0.483 | 1.596 | 0.53 |
| S-Realistic | 42662 | 21726 | 3727.8 | 0.058 | 0.460 | 2.692 | 0.55 |
| S-Flat | 21600 | 11020 | 1740.0 | 0.400 | 0.457 | 0.568 | 0.37 |
| S-Sine | 28950 | 14744 | 3439.0 | 0.301 | 0.573 | 0.929 | 0.53 |

The area of the simulated surface patch is 2350.4 mm^2^, which corresponds to about 1.3% of the whole cortical surface. The distance of the electrode contacts from the gray matter not included in the simulated (subcortical structures and excluded parts of the cortical surface) was between 5 and 13 mm, at least 3.5 times higher than the distance from the simulated part (Fig. 3C). The distance to the non-simulated parts is lower than the cut-off value 15 mm used to construct the surface due to the proximity of the hippocampus which was excluded by design.

On this surface the evolution of the system (1) was simulated. Since the model was supposed to capture only the activity on the cortical surface, the suspected source of the epileptic activity in the patient, right hippocampus, was not included in the simulation. Instead, noting the visible delay of the onset of the seizure activity on the laterally located contacts compared to mesially located contacts, we hand-placed the model epileptogenic zone to the mesial side of the extracted surface (Fig. 3B). This position approximately coincides with the right entorhinal cortex, to which is the right hippocampus strongly connected via the perforant path (Witter, 2007), supporting the hypothesis that the seizure spreads to the cortical patch from there. This model epileptogenic zone was represented as a small patch of diameter 5 mm with the value of excitability *u*_0_ increased above the critical level so that the tissue would start the seizure autonomously without any external intervention.

Parameters of the simulation (Tab. 1) were set as follows: The time constant of the whole system *τ*_*s*_ was set so that the frequency of the oscillations of the intermediate subsystem in an uncoupled oscillator at the seizure onset is the same as the dominant frequency in the SEEG recordings. Next, the time constant of the slow variable *τ*_0_ was set to the indicated value to obtain the seizure duration on the order of tens of seconds. Finally, we calculated the seizure spread velocity from the patient recording and performed a parameter sweep to find the strength of the fast-fast coupling *γ*_11_ which produces the same velocity. We assumed that the ictal wavefront travels in the direction of the electrode and we estimated the ictal propagation velocity from the temporal difference of the onsets on the first and last contacts and the geodesic distance of the surface points closest to these contacts. The resulting velocity was 5.98 mm s^−1^ which corresponds to the strength *γ*_11_ = 0.53 according to the results of the parameter sweep (Fig. S2).

### 3.2. Surrogate surfaces

In order to demonstrate the influence of the geometry of the cortical surface, we also performed the simulations of seizure spread on three surrogate surfaces: realistic, flat, and sine (Fig. 3B).

The realistic surrogate surface (S-Realistic) was constructed from the patient’s MRI the same way as the real surface, but using a different electrode. The electrode was also implanted in the right temporal lobe and was located posterior from the TB electrode, being embedded in the inferior temporal gyrus, fusiform gyrus, and parahippocampal gyrus. The flat surrogate surface (S-Flat) was constructed as a flat rectangular domain extending *m* = 15 mm on both ends of the electrode and in the direction perpendicular to the electrode orientation to both sides, thus having dimensions (*l* + 2*m*) × 2*m*, where *l* = 28 mm is the distance between the first and last contacts of the electrode. The surface was placed in the distance 1.47 mm from the electrode, so that the average distance from the contacts to the surface is the same as for the real surface. The sine surrogate surface (S-Sine) was constructed as a sine wave with wave length *l* and amplitude *A* = 7.89 mm selected so that the geodesic distance between the surface point closest to the first contact and the point closest to the last contact is the same as for the original surface. The extent of the sine surface is the same as of the flat surface.

Parameters of the triangulated surfaces are given in Tab. 2. For all three surfaces, the epileptogenic zone was placed close to the innermost contact of the electrode to reproduce the observed propagation pattern from the first to the last contact. As for the real surface, the coupling strength *γ*_11_ in the model was adjusted to produce the same temporal delay of the detected onset on the first and last contact pair as in the patient SEEG recording. The resulting values of *γ*_11_ are also given in Tab. 2.

### 3.3. Spatial scales of SEEG averaging

Fig. 3D shows the spatial plot of the gain matrix row associated with a single contact. The SEEG signal is generated by summing the weighted contributions of the source dynamics according to the values of the gain matrix row. To provide deeper insight into this forward model, Fig. 3E shows for what percentage of the gain matrix is a cortical area of given size responsible for. More specifically, we built the gain matrix for the full cortical area using the methods described in Sec. 2.5. For each contact (or contact pair in case of bipolar referencing) we then sorted the vertices by the absolute value of the gain matrix elements and plotted the accumulated absolute values of the gain matrix against the accumulated area. Simply put, the resulting curves show the spatial selectivity of all contacts or neighboring contact pairs. The fast saturation of the bipolar curves demonstrate the much higher spatial selectivity of the bipolar referencing compared to monopolar.

This notion is further elaborated by the figure inset which shows the size of the cortical area responsible for 50% of the gain. The forward model indicates that with monopolar referencing one needs around 30 cm^2^ of cortical surface to generate 50% of the SEEG signal. With bipolar referencing the size of the area drops below 1 cm^2^. While this is only a static view without taking into account the effect of the source dynamics, it provides important quantitative measure of the spatial selectivity of the implanted depth electrode.

### 3.4. Evolution of the simulated seizure

Let us have a look at the seizure spread simulated on the real surface, i.e. the surface around the investigated electrode. The seizure starts spontaneously in the small patch where the excitability was set to higher value (Fig. 4A, *t* = 5 s). From there it then spreads to the surrounding tissue (*t* = 10, 15 s), until the whole patch is recruited (*t* = 20 s). Around *t* = 30 s, the oscillatory activity starts to recede in the same direction as it spread before, and at *t* = 40 s the seizure activity has entirely terminated. The seizure does not terminate at once, even though the Epileptor field model (with appropriate parameter values) was shown to be able to support the synchronous termination (Proix et al., 2018). Considering that our main objective here was the investigation of the early seizure patterns, we have not searched the parameter space for the values where the simultaneous termination occurs.

**Figure 4:**
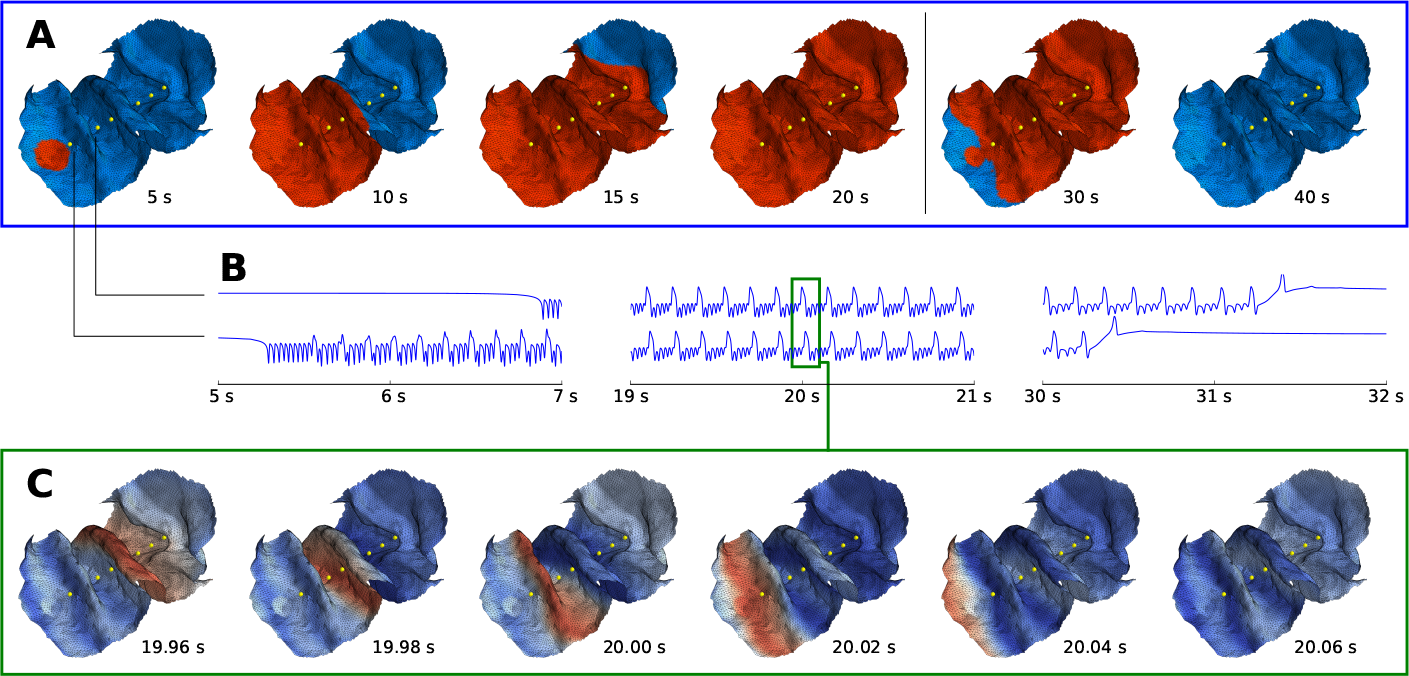
Evolution of the simulated seizure. (A) Extent of the oscillatory activity over time. Blue color represents the normal state (*u*_1_ < − 0.8) and red the seizure state (*u*_1_ ≥ −0.8). (B) Source activity *s*(*t*) in two points on the cortical surface shows the typical Epileptor dynamics with oscillations on two time scales. All points on the cortical surface follow qualitatively similar dynamics. (C) Snapshots of source activity *s*(*t*) over interval of 0.1 s during the seizure reveal the fast wave propagating across the cortical surface.

The source activity at any point on the cortical surface follows qualitatively the same dynamics typical for the Epileptor model (Fig. 4B): Sudden seizure onset as the fast subsystem of the Epileptor crosses the saddle-node bifurcation, then combination of the oscillations on two time scales during the seizure, and eventually the termination via homoclinic bifurcation.

Due to the local coupling of the Epileptor field, the oscillations of the intermediate system synchronize and form fast waves traveling across the cortical surface (Fig. 4C). Such waves were observed in the cortex of human patients during seizures, spatially extending not only across microelectrode arrays but also across ECoG arrays (Wagner et al., 2015; Smith et al., 2016; Martinet et al., 2017). In this simulation the velocity of the fast waves at time *t* = 20 s, when the whole patch was recruited, was estimated at 604 mm s^−1^, falling within the reported range of wave velocities of 100 to 1000 mm s^−1^ in human patients (Wagner et al., 2015; Smith et al., 2016; Martinet et al., 2017).

The simulated SEEG signals are plotted on Fig. 5 (cf. Fig. 1 with the real patient recording). We point out several features of the signals. Firstly, unlike the rapid onset on the source level, the amplitude of the SEEG signals grows gradually. The simulated signals therefore have the typical features of the TAA pattern, as in the real recording. The ramp-up phase, during which the amplitude grows, is on the order of several seconds, similarly as in the recording. Secondly, there is a clear temporal delay of the seizure onset on the lateral contacts, caused by the slow seizure spread. And lastly, the amplitude of the simulated signal varies across the contact pairs. Since the source dynamics is fairly stereotypical on the whole cortical patch, this is to be explained by the geometry of the surface and by the position of the contacts. We will analyze these features in the following section.

**Figure 5:**
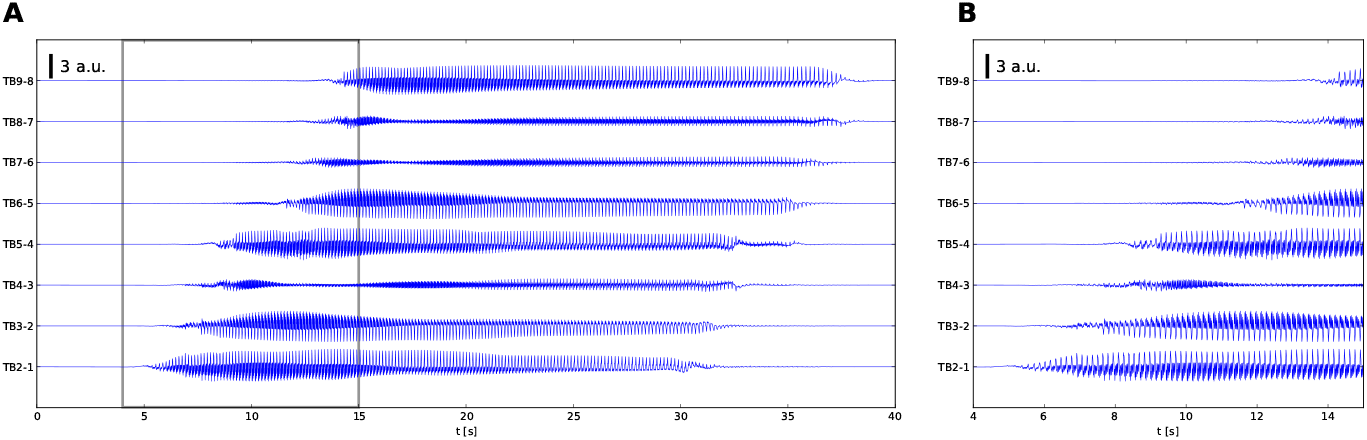
Traces of the simulated SEEG signals in bipolar representation. (A) Full simulated seizure. (B) Detail of the seizure onset, delimited by gray lines in (A).

### 3.5. Quantitative analysis

For quantitative analysis we identified the onset times and calculated the average amplitude of the simulated SEEG signals with bipolar referencing. Then we compared the values with the values obtained in the same way from the real SEEG recording (Fig. 6). The goodness of fit for all evaluated surfaces is shown in Tab. 3. Using the electrode-specific surface gives a better fit than the three surrogate surfaces in all evaluated criteria, supporting the necessity of using individual geometry.

**Figure 6:**
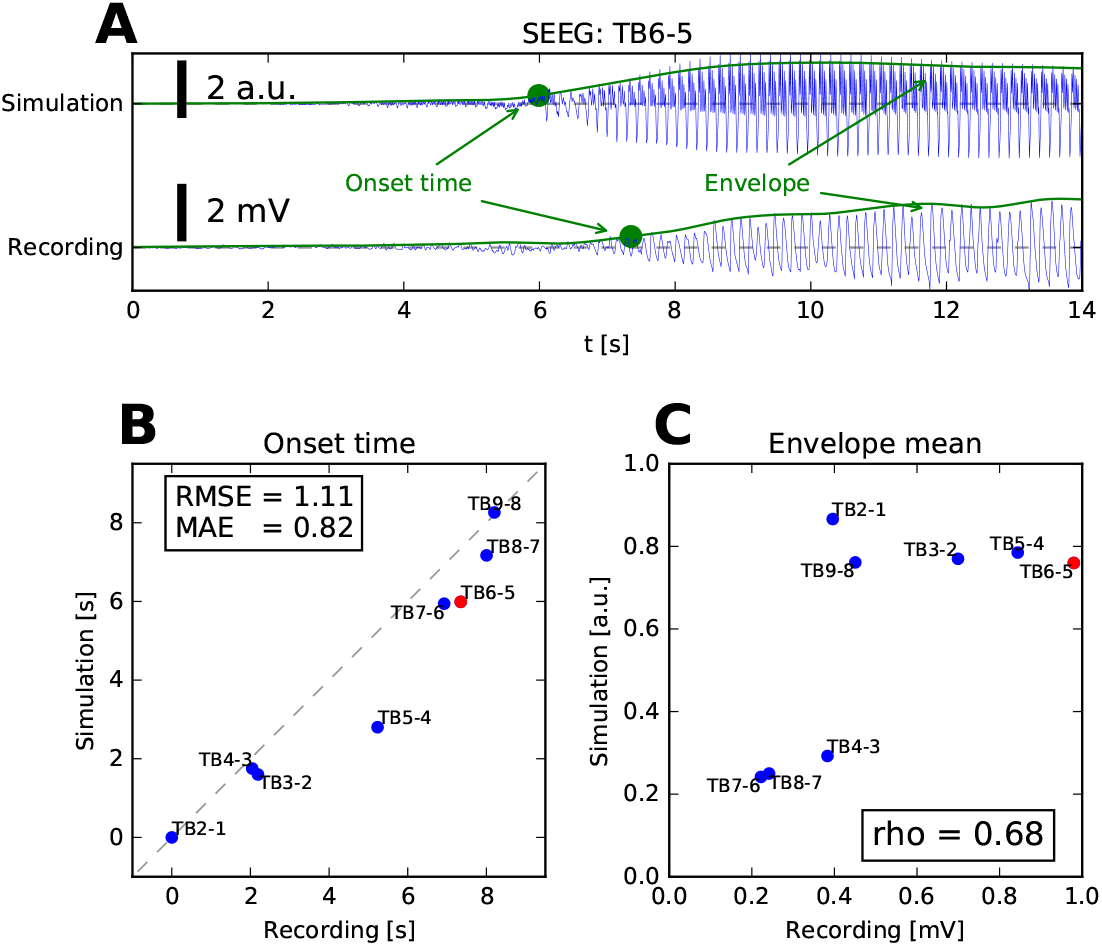
Quantitative analysis of the SEEG signals. (A) For all bipolar signals the envelope of the signal was calculated, and the onset time was determined (see Methods). The signals were shifted so that the onset time of the first bipolar signal was at zero. (B) Comparison of the onset times for the recorded SEEG and the SEEG simulated on the real surface. In the box two measures of fit are reported: root mean square error 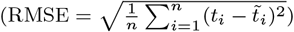 and mean absolute error 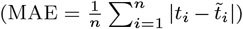. The red point corresponds to the signals shown in panel A. (C) Comparison of the mean value of the signal envelope for the recorded SEEG and the SEEG simulated on the real surface. The correlation coefficient *ρ* is reported in the box.

**Table 3:** Evaluation of the fit of the onset times and envelope means between the recorded SEEG and the SEEG simulated on the real and three surrogate surfaces. RMSE: root mean square error, MAE: mean absolute error, *ρ*: correlation coefficient (for definitions see the caption of Fig. 6). Best fit in each column is highlighted in bold.

| Surface | Onset time |  | Envelope mean |
| --- | --- | --- | --- |
| | RMSE | MAE | $\rho$ |
| Real | **1.11** | **0.82** | **0.68** |
| S-Realistic | 2.03 | 1.70 | 0.47 |
| S-Flat | 1.38 | 1.04 | 0.09 |
| S-Sine | 1.58 | 1.16 | -0.47 |

Looking at the amplitude of the signals, we can ask how much is this a result of the source dynamics and how much of the geometry of the cortical sources and position of the sensors. In our model, if the source dynamics of the whole surface patch was fully spatially homogeneous, i.e. all points were behaving exactly the same way, the amplitude of the bipolar signal between contacts *i* + 1 and *i* would be determined by the gain matrix *G* and would be proportionate to |∑_*j*_ *G*_*i*+*i,j*_ − *G*_*i,j*_|. Fig. 7 shows that this description is indeed close to the situation in the performed simulation of the seizure spread. The amplitude of the signals in the real recording can be also largely explained by the geometry of the sources and sensors.

**Figure 7:**
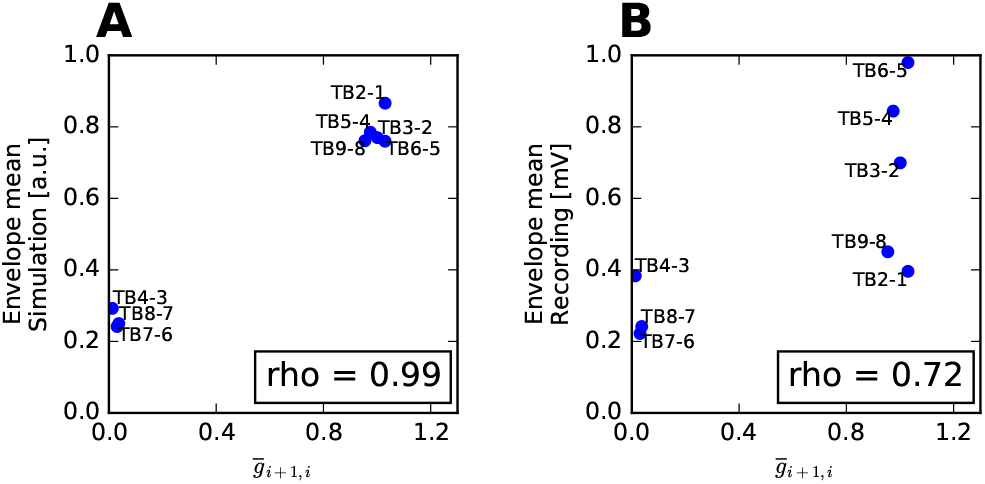
Influence of the gain matrix on the signal amplitude. (A) Correspondence between the term 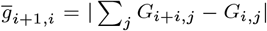 and the envelope mean of the simulated signals. The term 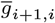 is the proportionality term between the source activity and the bipolar signal between (*i* + 1)th and *i*th contact assuming the source activity is spatially homogeneous on the whole cortical patch. Correlation coefficient close to one indicates that the amplitude of the simulated signals can be largely explained by the geometry of the sources and sensors. (B) Same as (A) for the recorded signals.

### 3.6. Spectral signatures of the simulated SEEG

Even though all SEEG signals arise from the stereotypical source activity, the geometric arrangement of the sources and sensors has a strong effect and the resulting signals might differ significantly. One specific example of this is illustrated on Fig. 8 which shows how the geometry can highlight or suppress activity in different frequency bands. Looking at the bipolar signals and their spectrograms only, one might classify the two patterns in different categories: TB6-5 shows the prominent 8 Hz oscillations with gradually increasing amplitude, and could be thus classified as the TAA pattern. However, the lower frequencies are suppressed on the neighboring TB7-6, where the activity in *γ* range is more prominent, so its features would point instead to the fast activity onset pattern. While such difference was not seen in the recorded SEEG (Fig. S3), our model indicates that the same source activity can manifest in different ways on the SEEG sensors, depending on the geometrical arrangement.

**Figure 8:**
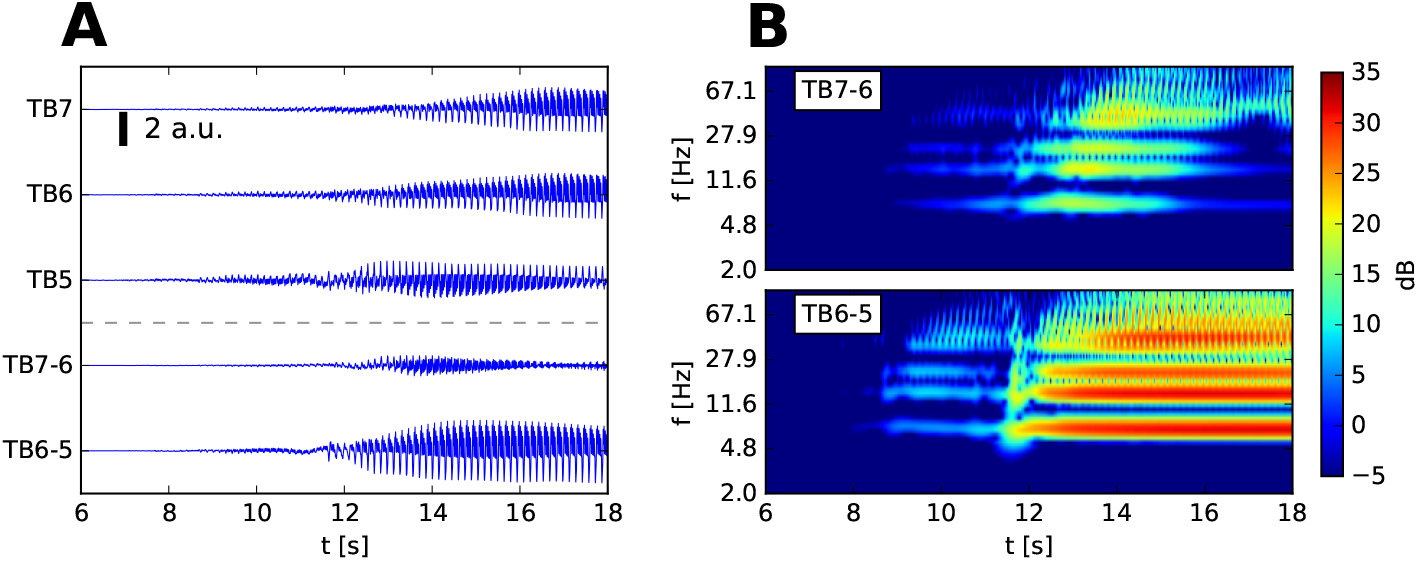
Geometry of the sources and sensors affects the spectral properties of the simulated SEEG signals. (A) Monopolar and bipolar view of seizure onset on selected contacts, as simulated on real surface. (B) Spectrogram of the two bipolar signals. 1/*f* normalization (spectral flattening) was applied to the spectrogram. The two bipolar signals differ notably in amplitude and its dominant frequencies. The low frequencies, present in TB6-5, are suppressed on TB7-6 due to the geometrical configuration of the cortical surface.

## 4. Discussion

### Main results

In this work we have investigated seizure front propagation across the cortical surface. We performed numerical simulations of the seizure spread on the cortical surface (i.e. in the source space) and from this source activity we generated the SEEG signals (i.e. in the sensor space) using the SEEG forward model. The results showed a qualitative and quantitative agreement with the patient recording: the TAA pattern was reproduced with similar duration of the ramp-up period of few seconds. The comparison with the simulations performed on the surrogate surfaces revealed that the real surface produces a better agreement with the recording in terms of the onset timing and the amplitude of the signals.

To obtain this good fit, three components were crucial: Firstly, it was the geometry of the cortical sources and position of the sensors in three-dimensional physical space, which we derived from the patient MRI scans. Secondly, it was the velocity of the seizure spread, which we estimated based on the onset delays in the SEEG recordings. The resulting value around 600 mm s^−1^ fell within the plausible range based on the previously published works. And lastly, it was the direction of the propagation, determined by the placement of the epileptogenic zone in the model. We have selected one characteristic patient and have demonstrated good qualitative and quantitative agreement with the patient SEEG recording. Our further analysis of various cortical surrogate surfaces including flat, sinusoidal and realistic, but not electrode-specific, surfaces demonstrate good sensitivity and specificity, thus promising usefulness in understanding the spatiotemporal organization of the seizure in clinical settings.

This work dealt with a single instance of TAA pattern, demonstrating that in this case there exists a plausible configuration of seizure origin, spreading direction, and spreading velocity that produces qualitatively similar SEEG features as those observed in the patient. This configuration was found by hand, and it is not guaranteed that it provides the best fit possible among all possible configurations. An approach which would find this optimal configuration for any given TAA instance and patient-specific geometry would be desirable and would provide further evidence for the relation between TAA and spreading seizures, however, given the large search space finding this solution may be computationally challenging.

### Classification of seizure onset patterns

Current approaches to the classification of the seizure onset patterns in clinical research rely on the analysis of frequencies, amplitudes, and waveforms present in the intracranial recordings (Doležalová et al., 2013; Perucca et al., 2014; Jiménez-Jiménez et al., 2015). However, as we have shown on Fig. 8, the same source dynamics of spreading seizure can produce electrographic seizure patterns from both TAA and fast activity classes. Curiously, these two patterns are the most common patterns in the areas of seizure propagation (Perucca et al., 2014). The results therefore raise the question if these patterns indeed reflect different source dynamics, or whether they are simply different manifestations of the same source dynamics caused by the complex geometry of the cortical sources.

In the example presented on Fig. 8 a careful examination of the SEEG signals with both bipolar and monopolar referencing would reveal the presence of low frequency oscillations and prevent the classification as the fast activity onset pattern. But two issues still remain: First, if the monopolar signals are dominated by the background noise or if the classification is done automatically based only on the features bipolar signal, the misclassification might not be easy to avoid. But more importantly, we argue that by not considering the geometry of the sources and sensors one leaves out an important piece of the puzzle. The SEEG signals are only a distorted reflection of the source activity, and it is the source activity and its dynamical patterns that should be analyzed and classified in order to gain deeper insight into the epileptogenesis. However, robust methods for the signal inversion from sensor to source space which would take into account the complex geometry still remain to be developed.

### Bifurcation-based classification

When analyzing the time series generated by an unknown biological process, one approach based in the nonlinear systems theory is to look for the sudden qualitative changes in behavior, such as the change from steady to oscillating state. From the features of this transition (such as changes in amplitude and frequency) one might infer the bifurcation that the underlying dynamical system is going through during this transition. The equations of any dynamical system undergoing a specific bifurcation can be smoothly mapped to the so-called normal form, which conserve the qualitative behaviour of the system around the bifurcation. The knowledge of the bifurcation type that the system is going through might then guide the development of a mathematical model of the observed activity (Kuznetsov, 1998; Izhikevich, 2010; Touboul et al., 2011; Jirsa et al., 2014). A systematic analysis of onset and offset bifurcations then permits the construction of a seizure taxonomy based on purely dynamic features (Saggio et al., 2017, 2020).

From this perspective the TAA pattern would correspond to a dynamical system undergoing a supercritical Hopf bifurcation, whose distinguishing feature is the gradually increasing amplitude of the oscillations after crossing the bifurcation (Izhikevich, 2010). However, as we have shown in this paper, the features that are seen on the generated SEEG signals might not be present on the source level. Instead of a Hopf bifurcation, our employed model of the source dynamics switches to the seizure state by crossing the saddle-node bifurcation. The type of the onset bifurcation has important consequences for the behavior of the dynamical system. One of them is the response to external stimulation. A monostable system going through a supercritical Hopf bifurcation transitions from a fixed point to a limit cycle. Before the transition, a brief stimulation would not be able to force the monostable system to a sustained oscillatory state, since there exits only a single fixed point. On the other hand, a system undergoing a saddle-node bifurcation might be sensitive to stimulation: before the transition the system resides in a bistable state with a stable fixed point and stable limit cycle and stimulation of the system might therefore move it to the other stable state. This difference between the two dynamical systems highlights the importance of performing the bifurcation analysis on the source level, pointing us again to the need for robust sensor-source inversion methods.

## Author contributions

V.S. and V.J. designed the study. M.G. and F.B. acquired the data. V.S. performed the study. V.S., M.G, F.B., and V.J. wrote the manuscript.

## Acknowledgements

The authors wish to acknowledge the financial support of the following agencies: the French National Research Agency (ANR) as part of the second “Investissements d’Avenir” program (ANR-17-RHUS-0004, EPINOV), European Union’s Horizon 2020 Framework Programme for Research and Innovation under the Specific Grant Agreement No. 785907, PHRC-I 2013 EPISODIUM (grant number 2014-27), the Fondation pour la Recherche Médicale (DIC20161236442) and the SATT Sud-Est (827-SA-16-UAM) for providing funding for this research project.

## Data availability

The data set cannot be made publicly available due to the data protection concerns, but can be made available upon reasonable request to authors.

## Code availability

The code used in the study will be made available upon publication.

## Supplementary material

**Figure S1:**
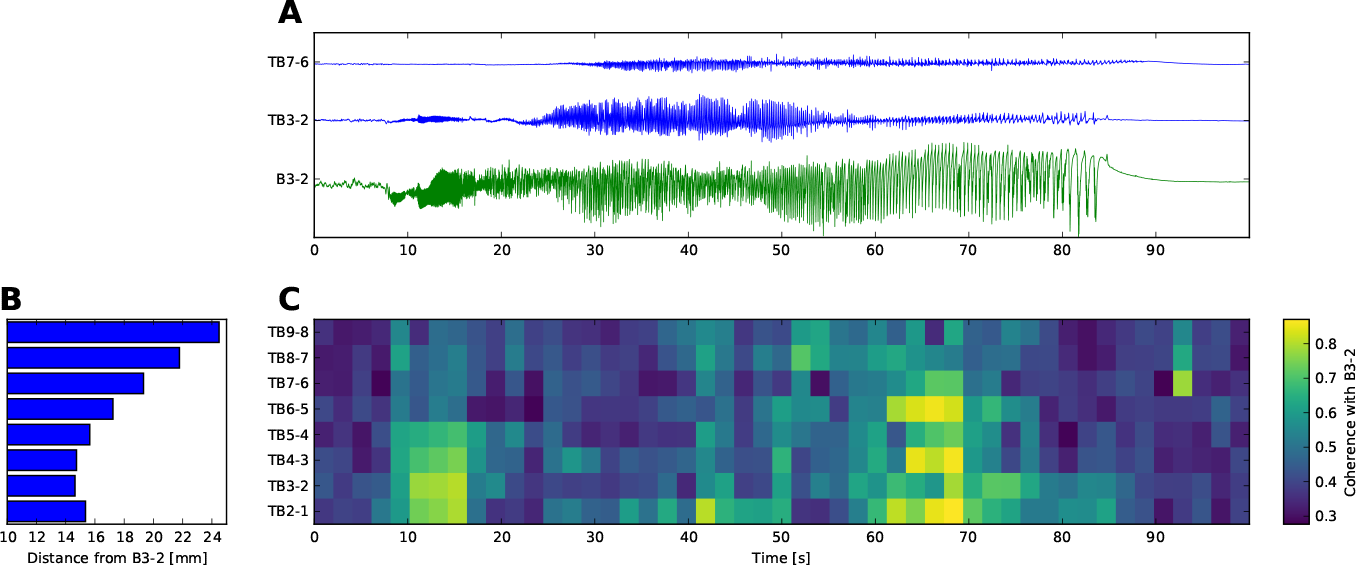
Coherence indicates that the initial oscillatory activity on the contacts of the TB electrode are due to the volume conduction. (A) Selected bipolar traces. On B3-2 and TB3-2, a fast (∼20 Hz) oscillatory activity appears between 8 and 18 s, with high amplitudes at B3-2 and lower at TB3-2. (B) Euclidean distance of a midpoint between two neighboring contacts to the midpoint B3-2. (C) Average coherence with B3-2 in a frequency band [1, 30] Hz. At 8 s, high coherence values are appears on channels TB2-1 to TB5-4, disappearing again after 18 s. In this period the coherence is highest at TB3-2, which is nearest to B3-2, and decays with increasing distance. High values of coherence reappear later (60 - 70 s) as the seizure progresses, however, without the clear dependence on the distance from B3-2. In our view, this indicates that the initial oscillations on all electrodes reflect the activity at right hippocampus, where B3-2 is placed, and are seen only due to the volume conduction. The later oscillations on the other hand reflect the source activity close to the contacts, and are coherent due to the source synchronization.

**Figure S2:**
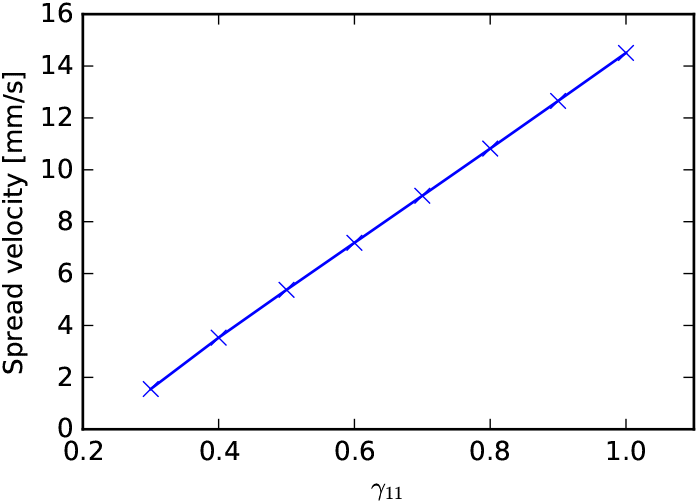
Dependence of the seizure spread velocity on the coupling strength *γ*_11_. The relation was determined by numerical simulations of a seizure spread on a 2D rectangular patch with varying *γ*_11_ (see Methods). For *γ*_11_ < 0.3 the seizure failed to propagate from the epileptogenic zone.

**Figure S3:**
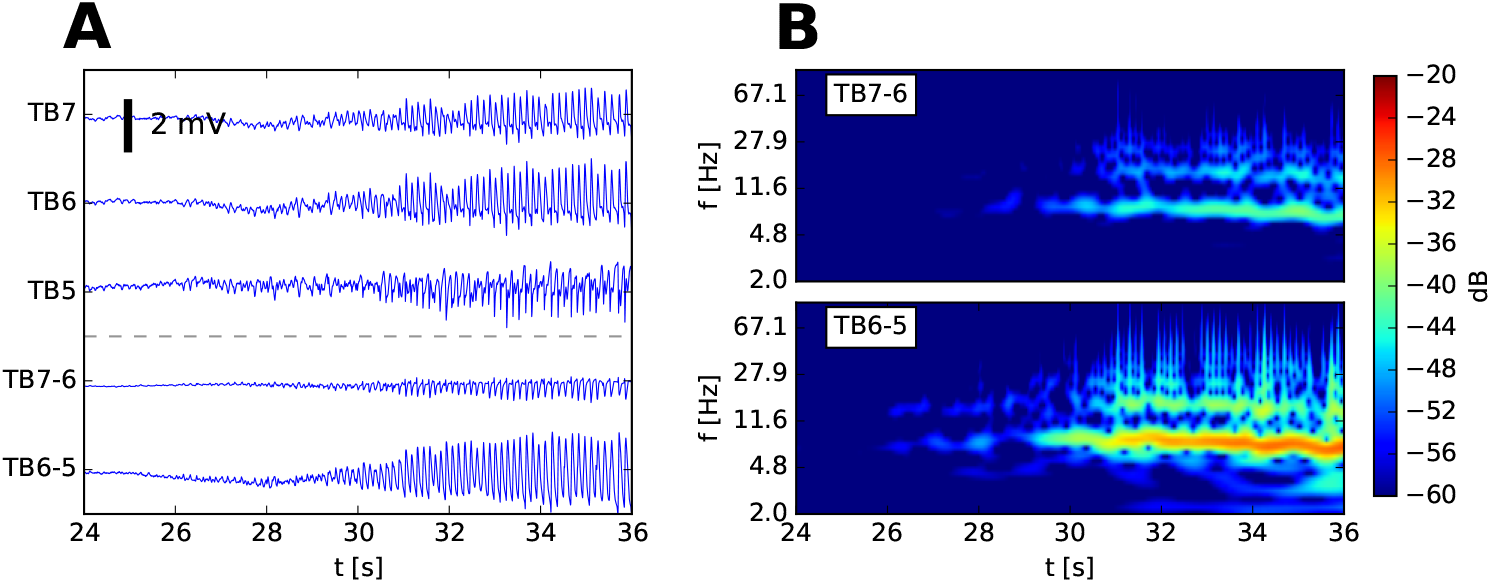
Spectral signatures of the recorded SEEG signals. Cf. Fig. 8. (A) Monopolar and bipolar view of seizure onset on selected contacts. (B) Spectrogram of the two bipolar signals. 1/*f* normalization (spectral flattening) was applied to the spectrogram.

